## Supplementary material for "Tracking H3K27me3 and H4K20me1 during XCI reveals similarities in enrichment dynamics": All supplemental files: Figure EV1.pdf

Fig. EV1

**A**

4 MAQVQLQQSGAELVKPGASVKLSCKASGFTFTSYYMYWVKQRPGQGLEWIGEINPSNGGT

$V_H$

86

64 YFNEKFKNKATLTVDKSSSTAYMHLSSLTSEDSAVYYCAAYYGNLFDYWGQGTTLTVSSG

158

124 GGGSGGGGSGGGGSDIVMSQSPSSLAVSVGEKVTMACKSSQSLLYSSNQKNYLAWYQQKP

$V_L$

184 GQSPKLLIYWASTRESGVPDRFTGSGSGTDFTLTISSVKAEDLAVYYCQQYYTYPWTFGG

244 GTKLEI

**B**

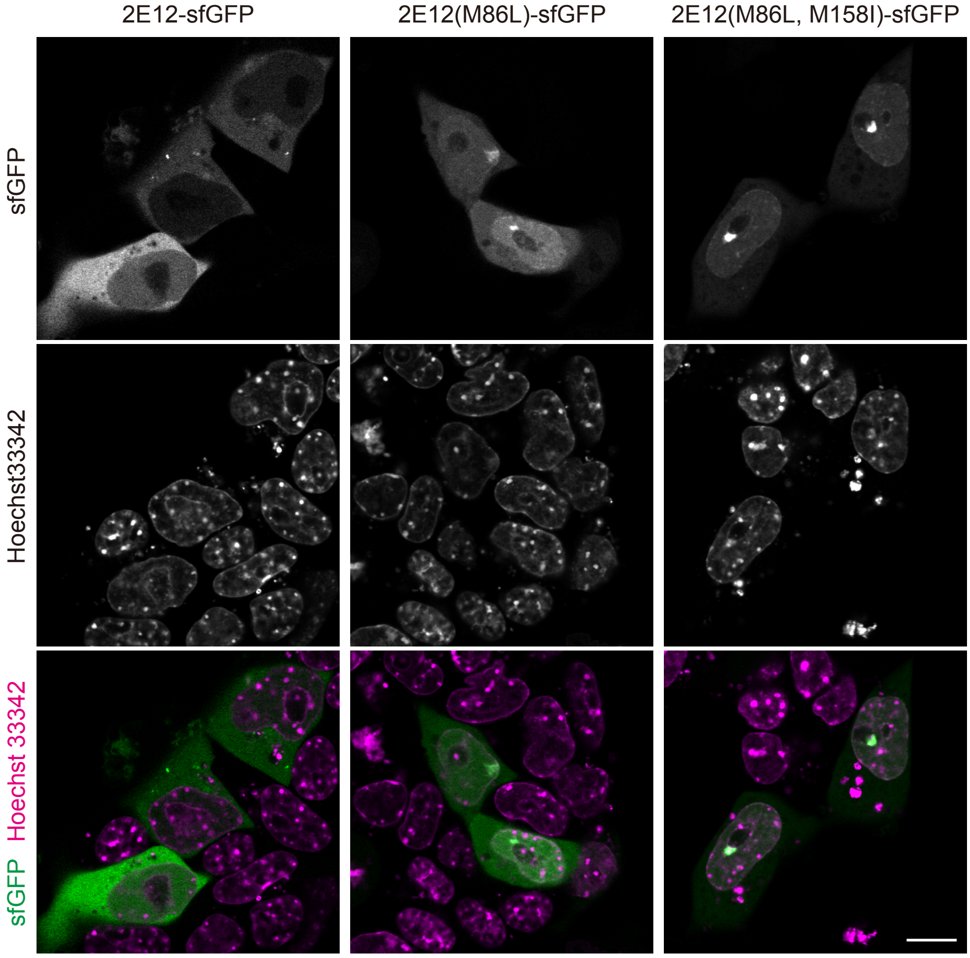


**C**

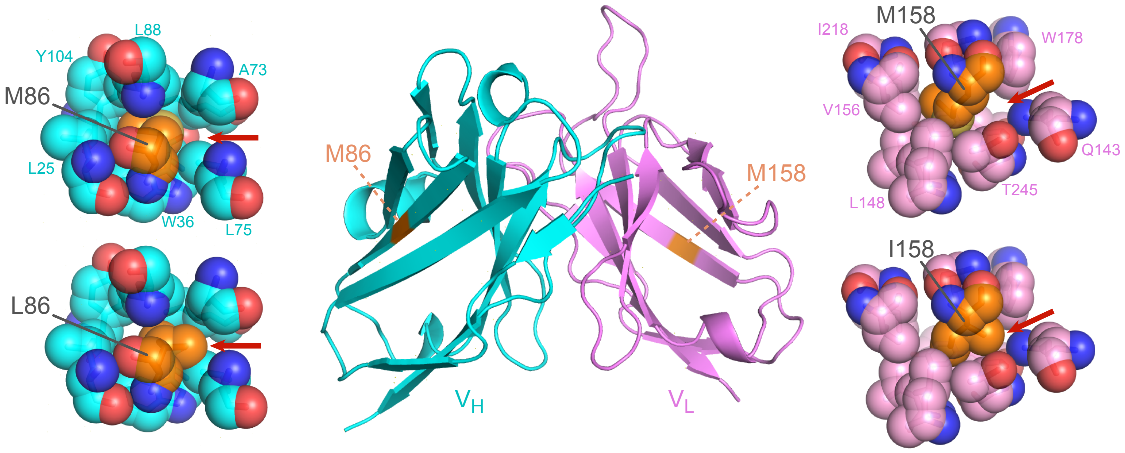
