## Supplementary material for "Tracking H3K27me3 and H4K20me1 during XCI reveals similarities in enrichment dynamics": All supplemental files: Figure EV2.pdf

Fig. EV2

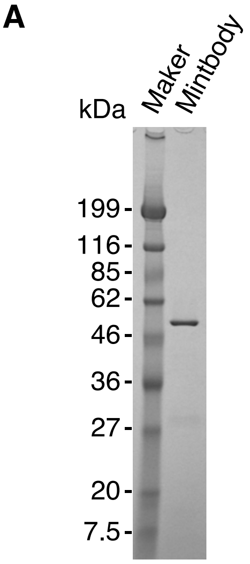


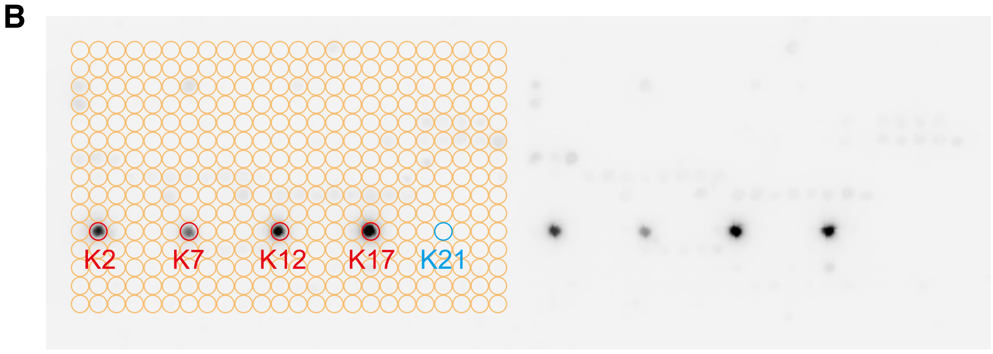


| Peptide location | Peptide sequence | Mod1 | Mod2 |
| --- | --- | --- | --- |
| K2 | PRKQLATKAARKme3SAPATGG | K27me3 | |
| K7 | PRKQLATKAARme2sKme3SAPATGG | R26me2s | K27me3 |
| K12 | PRKQLATKAARme2aKme3SAPATGG | R26me2a | K27me3 |
| K17 | PRKQLATKAARcitKme3SAPATGG | R26cit | K27me3 |
| K21 | PRKQLATKAARKme3SphAPATGG | K27me3 | S28ph |

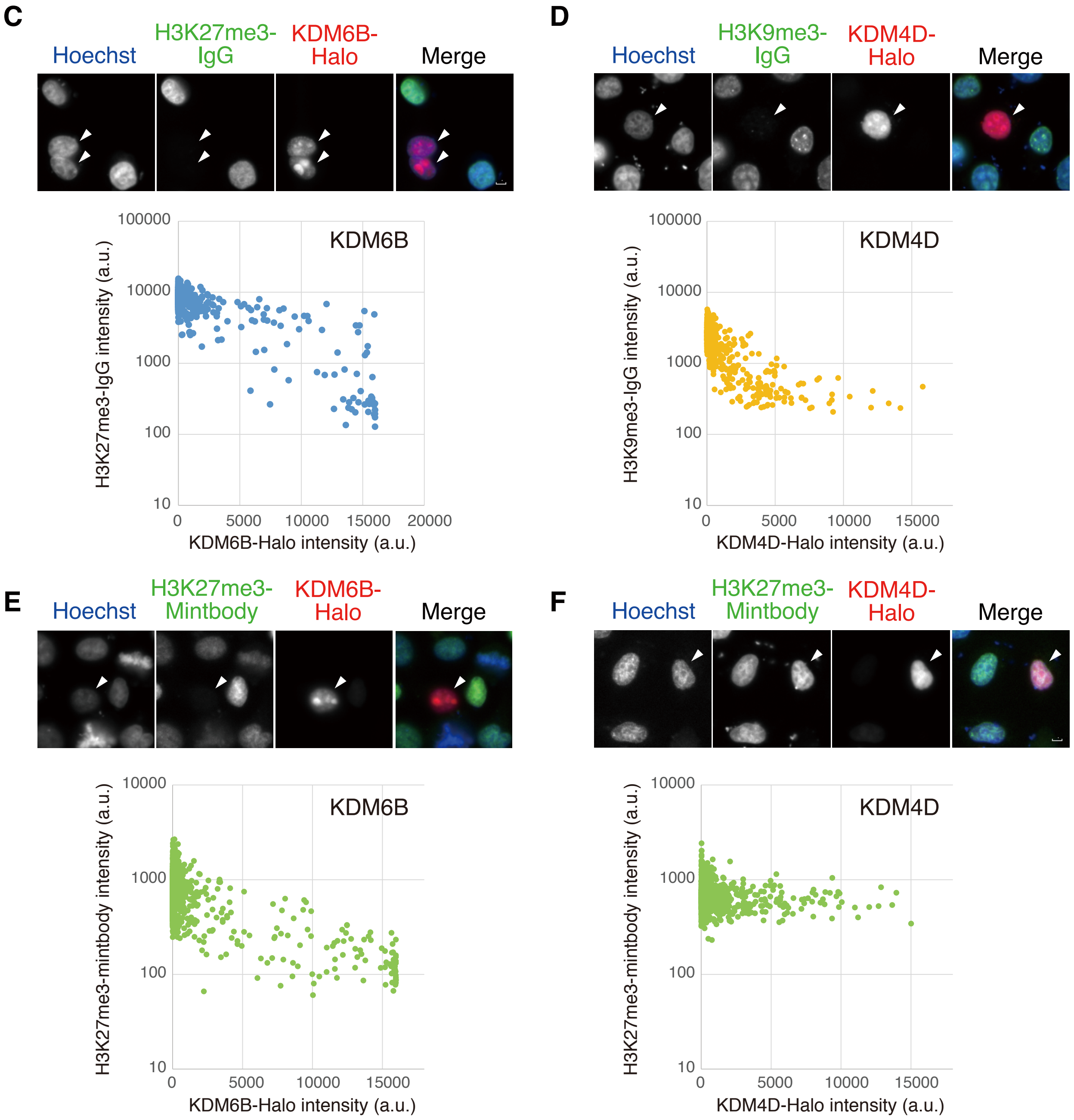
