## Supplementary material for "Tracking H3K27me3 and H4K20me1 during XCI reveals similarities in enrichment dynamics": All supplemental files: Figure EV5.pdf

Fig. EV5

A

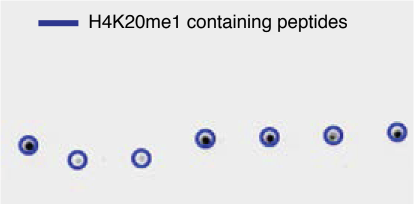


| Peptide location | Peptide sequence | Mod1 | Mod2 | Mod3 |
|---|---|---|---|---|
| N 1 | G K G G A K R H R Kme1 V L R D N I Q G I T | **K20me1** | | |
| N12 | G K G G A Kac R H R Kme1 V L R D N I Q G I T | K16ac | **K20me1** | |
| N16 | G Kac G G A Kac R H R Kme1 V L R D N I Q G I T | K12ac | K16ac | **K20me1** |
| N20 | G K G G A K R H Rme2a Kme1 V L R D N I Q G I T | R19me2a | **K20me1** | |
| N24 | G K G G A K R H Rme2s Kme1 V L R D N I Q G I T | R19me2s | **K20me1** | |
| O 4 | G K G G A K R H R Kme1 V L Rme2a D N I Q G I T | R24me2a | **K20me1** | |
| O 8 | G K G G A K R H R Kme1 V L Rme2s D N I Q G I T | R24me2s | **K20me1** | |

B

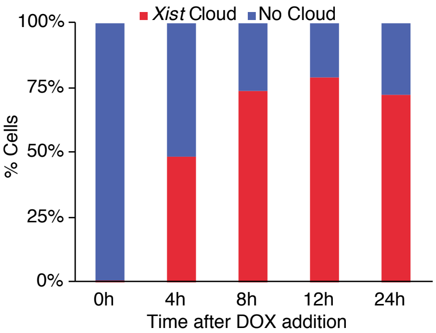


C

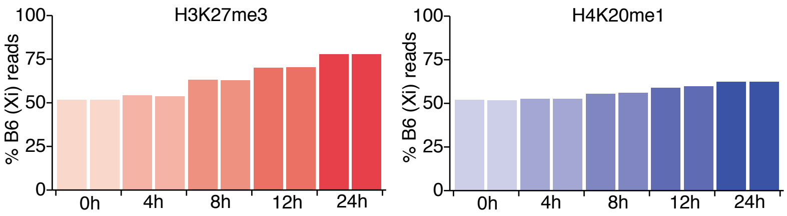


D

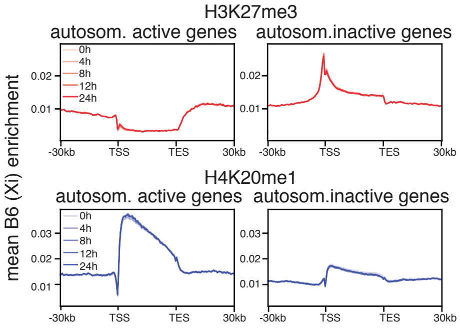


E

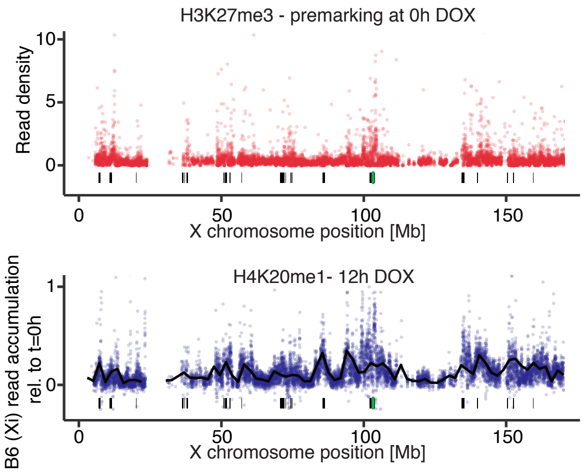
