## Supplementary figures and images for "Tracking H3K27me3 and H4K20me1 during XCI reveals similarities in enrichment dynamics"

### Figure EV3.pdf

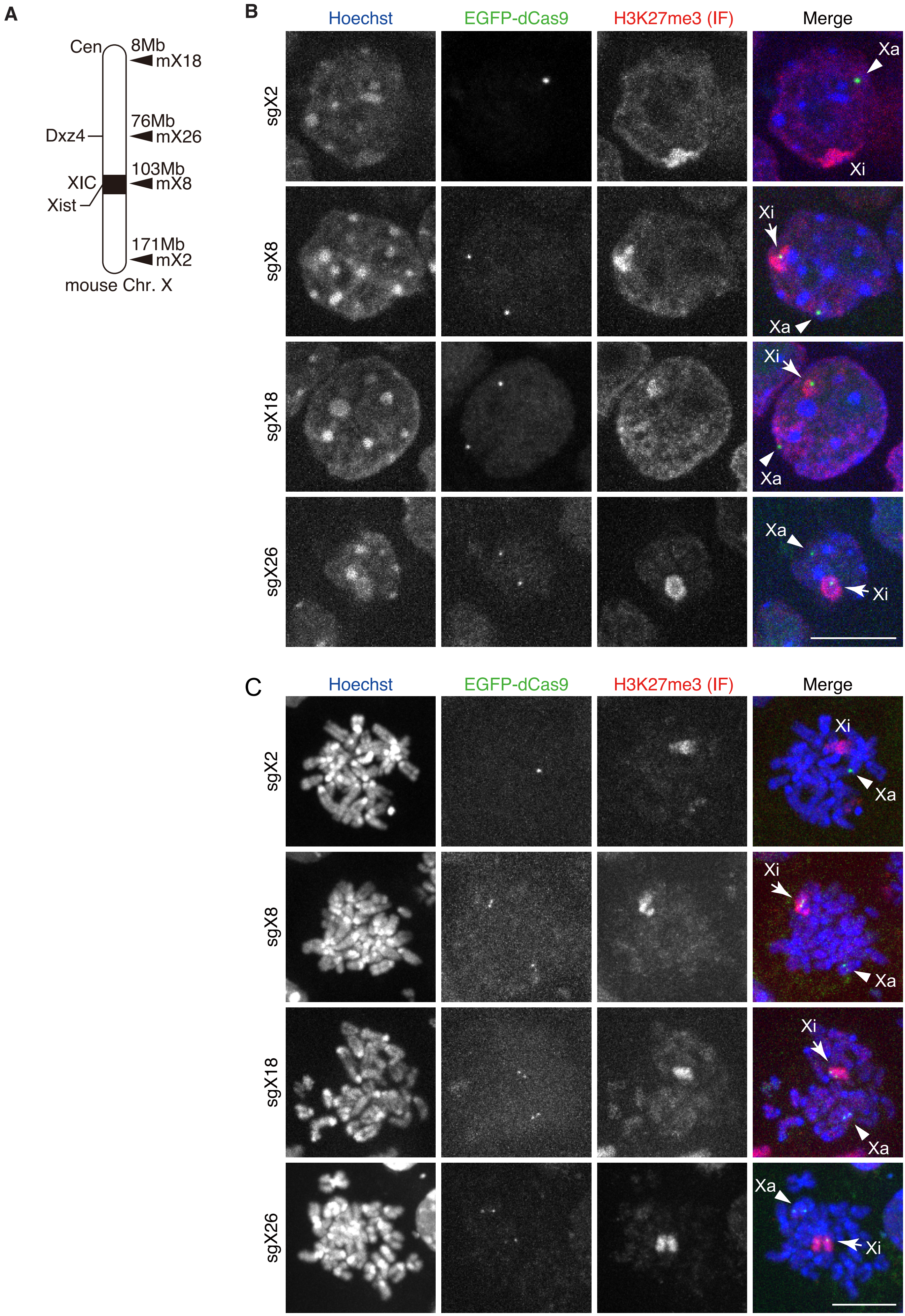

### Figure EV4.pdf

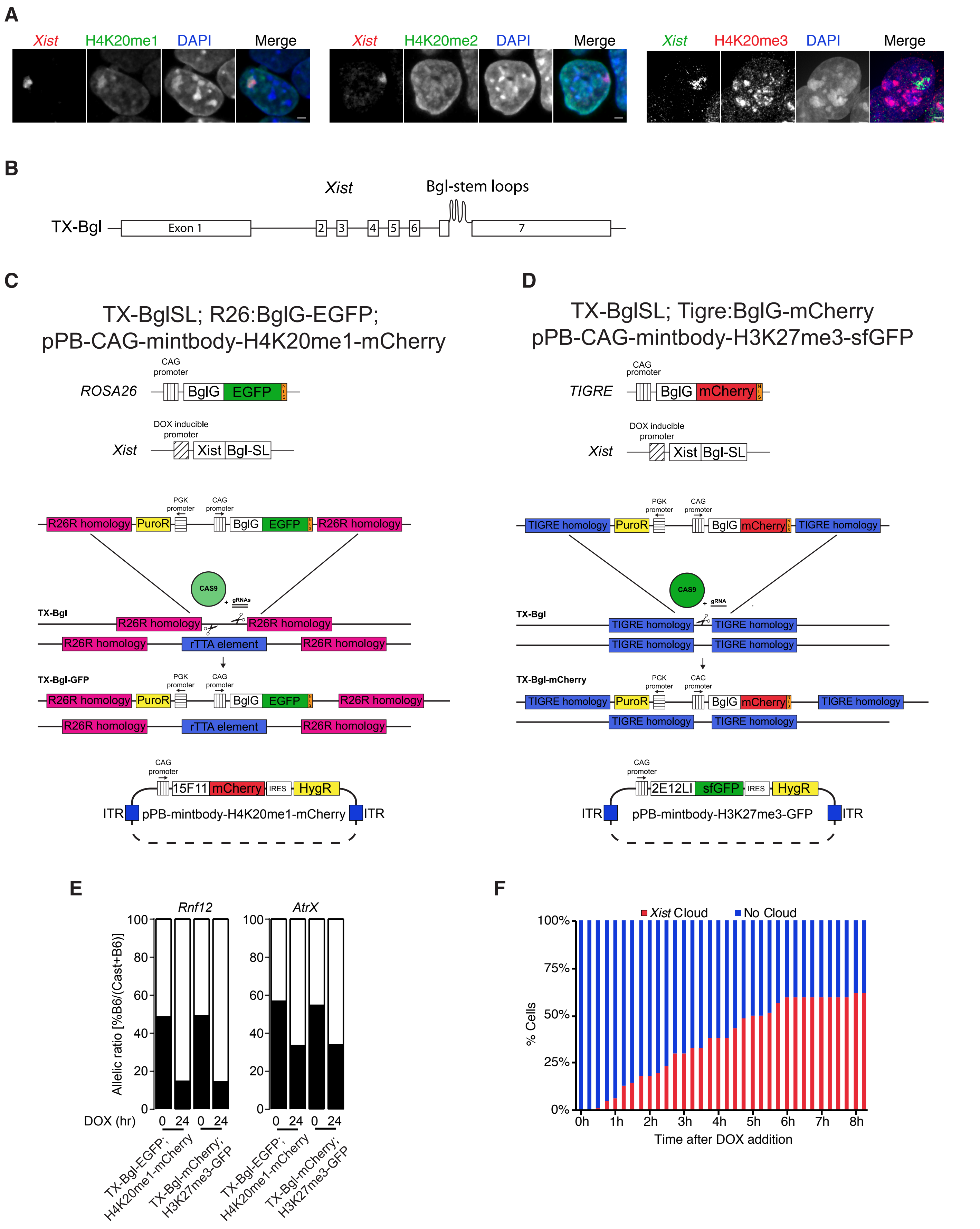
